# *In Silico* identification and modelling of FDA-approved drugs targeting T-type calcium channels

**DOI:** 10.1101/2024.09.27.615366

**Authors:** Pedro Fong, Susana Roman Garcia, Melanie I Stefan, David C Sterratt

**Author notes:** Corresponding authors: (PF), (DS).

## Abstract

Studies have shown that inhibition of the Ca_v_3.1 T-type calcium channel can prevent or suppress neurological diseases, such as epileptic seizures and diabetic neuropathy. In this study we aimed to use *in silico* simulations to identify a U.S. Food and Drug Administration (FDA)-approved drug that can bind to the Ca_v_3.1 T-type calcium channel. We used the automated docking suite GOLD v5.5 with the genetic algorithm to simulate molecular docking and predict the protein-ligand binding modes, and the ChemPLP empirical scoring function to estimate the binding affinities of 2,115 FDA-approved drugs to the human Ca_v_3.1 channel. Drugs with high binding affinity and appropriate pharmacodynamic and pharmacokinetic properties were selected for molecular mechanics Poisson–Boltzmann surface area (MMPBSA) and molecular mechanics generalised Born surface area (MMGBSA) binding free energy calculations, GROMACS molecular dynamics (MD) simulations and Monte Carlo Cell (MCell) simulations. The docking results indicated that the FDA-approved drug montelukast has a high binding affinity to Ca_v_3.1, and data from the literature suggested that montelukast has the appropriate drug-like properties to cross the human blood-brain barrier and reach synapses in the central nervous system. MMPBSA, MMGBSA and MD simulations showed the high stability of the montelukast-Ca_v_3.1 complex. MCell simulations indicated that the blockage of Ca_v_3.1 by montelukast reduced the number of synaptic vesicles being released from the pre-synaptic region to the synaptic cleft, which may reduce the probability and amplitude of postsynaptic potentials.

**Author Summary:** New drugs for current illnesses or disorders are always necessary to enhance therapeutic outcomes or can be utilised for patients who have experienced side effects from current medicines. The Ca_v_3.1 T-type calcium channel has been found to be linked to neurological diseases, such as epilepsy and diabetes. Consequently, identifying a drug that could bind to these receptors may yield therapeutic effects on these neurological conditions. This study utilises various computational methods to sift through 2,115 FDA-approved drugs that have been used for other diseases, not neurological ones, to determine whether some of these drugs could bind to the Ca_v_3.1 T-type calcium channel and potentially produce therapeutic effects on neurological diseases. Our research indicates that the current asthma drug montelukast has the potential to bind to the calcium channel and is worthy of further investigation.

## Introduction

Calcium channels are located in the plasma membrane of excitable cells, including neurons, brain cells and heart muscle cells [1]. These channels allow the influx of calcium ions into the cells and cause depolarisation and excitation, contributing to the biological functions of the cells [1]. Blocking the influx of calcium ions has been shown to have pharmacological effects. For example, ziconotide is a calcium channel blocker medication that inhibits the influx of calcium ions into neurons, consequently reducing neurotransmitter release and dampening neuronal excitability. This reduction in excitability leads to its analgesic effects and makes ziconotide a treatment for severe chronic pain [2,3].

Calcium channels can be categorized into voltage-gated and ligand-gated; the opening and closing of the former responds to a voltage difference, while the latter is governed by ligand binding. Voltage-gated calcium channels are subcategorised according to their response to voltage and temporal dynamics into L-type (long-lasting), N-type (neural), P-type (Purkinje), R-type (residual) and T-type (transient) [4]. Their distribution throughout the human body varies by type; for example, N-type channels are typically found in the brain, R-type channels in neurons, and L-type channels in muscles, bones, myocytes and dendrites. This study aims to find an inhibitor of the T-type channel, which is widely distributed across the central nervous system [4].

T-type calcium channels have three subcategories: Ca_v_3.1, Ca_v_3.2 and Ca_v_3.3, which are all associated with neurological diseases, such as epilepsy, neuralgia, diabetic neuropathy, nerve injury and sleep disorders [5]. Ca_v_3.1 is primarily expressed in the central nervous system and is involved in the generation of rhythmic activities such as sleep patterns and regulating neuronal firing patterns, and its over-activation may cause neurons to fire at an abnormally high frequency leading to epileptic seizures [5]. A prior study provoked seizures in wild-type mice lacking Ca_v_3.1 through intraperitoneal drug administration, revealing that the Ca_v_3.1 knockout mice have significantly less risk of absence seizures [4]. Ca_v_3.2 contributes to various physiological processes such as hormone secretion, neurotransmitter release, and regulation of cardiac pacemaker activity [6]. It is also associated with epilepsy, but its activation may not be strong enough to produce seizures [6]. Ca_v_3.3 channels are mainly found in the thalamus and play a crucial role in generating and modulating neuronal oscillations, sensory processing, and serving as the primary pacemaker for sleep spindles [7]. Thus, all three types of calcium channels are considered potential drug targets for neurological diseases [1]. Ca_v_3.1 is the most studied and is most associated with epileptic seizures. The objective of this study is to identify a U.S. Food and Drug Administration (FDA)-approved drug that can bind to the Ca_v_3.1 T-type calcium channels using *in silico* simulations.

The first stage of most drug discovery projects is *in silico* simulations [8], which help to identify potential drug candidates for further experiments. In general, the process starts with the creation of a library containing a large number of molecules. This library can be obtained from a chemical compound database, such as the ZINC database [9]. ZINC classifies compounds into different categories; for example, herbal ingredients, human metabolites and FDA-approved drugs. After the selection and compilation of a compound library, screening is performed to identify potential molecules that can bind to the selected therapeutic target. The drug-like properties, such as adsorption, distribution, metabolism, excretion and toxicity, of these compounds are then predicted using various *in silico* methods [10]. If required, the structures of these molecules can be modified in order to bind more tightly to the target or to have advanced drug-like properties, such as fewer side effects and higher bioavailability.

The *in silico* methods used in this study were molecular docking, molecular mechanics Poisson–Boltzmann/generalised Born surface area (MMPBSA/MMGBSA), molecular dynamics (MD) and Monte Carlo (MCell) simulations. The docking simulations were employed to identify an FDA-approved drug that has a high potential to bind with Ca_v_3.1. The MMPBSA/MMGBSA and MD simulations were used to validate that the identified FDA-approved drug can bind to Ca_v_3.1 with high stability. A pre-synaptic MCell model was created to simulate the effect of the chosen FDA-approved drug on lowering neurotransmitter release and weakening the neural signal by competing with calcium ions for the Ca_v_3.1 channels.

Using these methods, we identified montelukast as a promising candidate to bind Ca_v_3.1 channels and reduce the hyperactivity of presynaptic neurons. Further experimental studies are necessary to establish the feasibility of repurposing montelukast for the treatment of epilepsy and related conditions.

## Results and Discussion

### Molecular docking

Docking simulations were performed between the selected ligand-bound Ca_v_3.1 structure (Fig 1, PDB code 6KZP) and 2,115 FDA-approved drugs. The results show that 234 drugs obtained a higher binding score than the native ligand embedded in the crystal structure of the Ca_v_3.1 (PDB: Z944), indicating that these compounds may inhibit opening of the calcium channel and block the entry of calcium ions into the neurons. The FDA-approved drugs with the top 20 binding scores are shown in Table 1. Indocyanine green obtained the highest score. Its 2D chemical structure is larger than the Z944 structure and contains more flexible single bonds, which are located near to the two highly polar bisulphite groups (Fig 2). These features may enable the indocyanine green to fit more naturally in the binding site of Ca_v_3.1. The second (cobicistat) and third (ritonavir) ranked drugs also have larger and more flexible structures than the native ligand (Fig 2).

**Fig 1.**
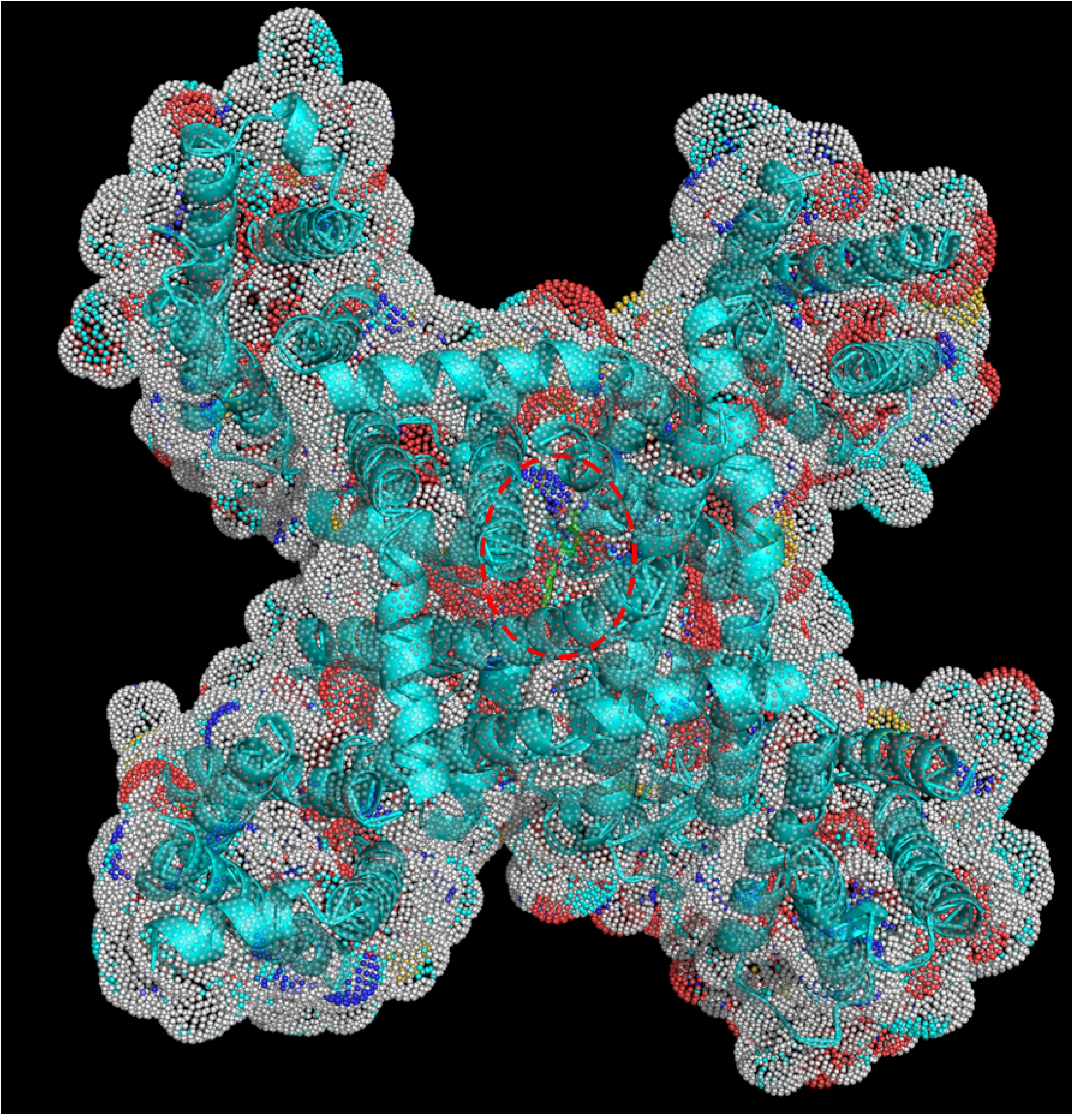
Surface and cartoon view of the native ligand (Z944) in the human Ca_v_3.1 (PDB: 6KZP). The binding site is indicated by the red dotted line.

**Fig 2.**
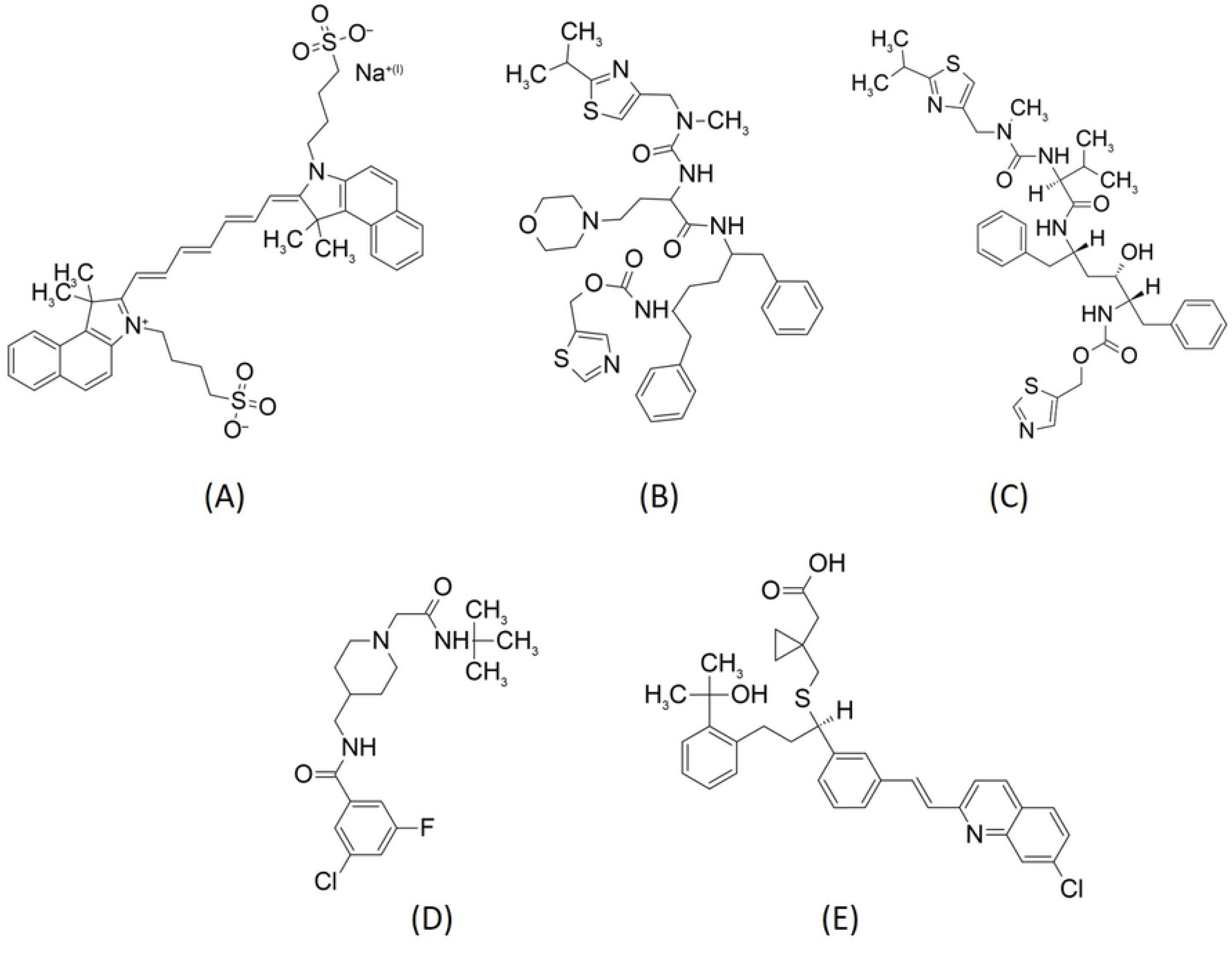
Chemical structures of (A) indocyanine green, (B) cobicistat, (C) ritonavir, (D) native ligand (PDB: 6KZP) and (E) montelukast.

**Table 1.** Clinical uses of the top 20 docking scored FDA-approved drugs.

| Drug name | Score | Clinical use |
| --- | --- | --- |
| Native (Z944) | 62.90 | No clinical usage |
| Indocyanine green | 110.68 | Diagnostic agent uses in angiography |
| Cobicistat | 110.63 | CYP3A inhibitor uses to increase the efficiency of<br>antivirus |
| Ritonavir | 108.20 | HIV protease inhibitors |
| Glycerol phenylbutyrate | 107.56 | Uses for Inborn urea cycle disorders |
| Carfilzomib | 102.35 | Proteasome inhibitors used in multiple myeloma |
| Saquinavir | 101.96 | HIV protease inhibitors |
| Dabigatran | 101.49 | Uses as anticoagulant |
| Montelukast | 100.71 | Leukotriene inhibitor use for asthma |
| Fosinopril | 100.48 | Angiotensin-converting enzyme inhibitor uses<br>for<br>hypertension |
| Dronedarone | 100.44 | Antiarrhythmic drug |
| Azilsartan | 99.26 | Angiotensin II receptor antagonist uses for<br>hypertension |
| Gadofosveset | 98.86 | Gadolinium-based MRI contrast agent |
| Atorvastatin | 98.84 | Statin uses for hypercholesterolemia |
| Naloxegol | 97.43 | μ-opioid receptor antagonist uses for constipation |
| Indinavir | 96.74 | HIV protease inhibitor |
| Silodosin | 96.64 | $\alpha_1$-adrenoceptor antagonist uses for benign pro-<br>static hyperplasia |
| Nonoxynol-9 | 96.42 | Vaginal spermicide |
| Isavuconazonium | 96.23 | Triazole uses as antifungal agent |
| Lomitapide | 95.40 | Microsomal triglyceride transfer protein inhibitor uses for hypercholesterolemia |

Although indocyanine green obtained the highest score, we do not believe that it is suitable for the drug repurposing aim of this study. We aimed to find a Ca_v_3.1 inhibitor to treat neurological disorders, such as epileptic seizures, that generally require long-term regular use. However, indocyanine green is a diagnostic agent for angiography, and no studies can be found on the safety of its long-term regular use [11]. Thus, regular administration may not be appropriate. Cobicistat obtained the second highest binding score (Table 1); it is a CYP3A (cytochrome P450, family 3, subfamily A) inhibitor used to increase the systemic exposure and hence the effectiveness of the HIV drugs atazanavir or darunavir [12]. CYP3A is a group of enzymes involved in the metabolism of many drugs; thus, cobicistat can cause various drug-drug interactions [12], and we do not therefore believe it to be appropriate for regular use.

The third- and fourth-ranked drugs were ritonavir and saquinavir. They are HIV antiviral drugs, and using them regularly may significantly increase drug resistance and worsen the global problems of HIV drug effectiveness [13]. Glycerol phenylbutyrate was the fifth-ranked drug; it is an oral medication for inborn urea cycle disorders. However, it has a poor side effect profile: 16% of patients had diarrhoea, 14% had flatulence, 14% had a headache, and 7% had abdominal pain [14]. The next ranked drug is carfilzomib, which is used as a treatment for myeloma. It only has parenteral administration and can cause serious side effects [15]. Dabigatran is the next ranked drug; it is an anticoagulant with a high risk of bleeding as a side effect [16]. Thus, all the above-mentioned drugs may not be appropriate for regular use to treat neurological disorders, such as epileptic seizures.

Montelukast obtained the eighth highest ranking score of 100.7 (Table 1), which is higher than the value of 62.9 for Z944. The docking conformation analysis shows that there were 15 hydrophobic contacts between montelukast and the amino acid residues in the binding site (Figs 3A and B). In contrast, the native ligand had hydrophobic contact with only 10 residues (Figs 3C and D). Most of these residues are different for montelukast and Z944; only three, Leu 872, Leu 920 and Phe 956, possess hydrophobic contacts with both montelukast and Z944. The other contact residues for montelukast are Ile 351, Thr 352, Leu 353, Ser 383, Ile 387, Gln 922 and Tyr 953 for Z994, and Asn 388, Leu 391, Phe 917, Thr 921, Asn 952, Lys 1462, Val 1505, Leu 1506, Phe 1509, Gln 1816, Val 1820 and Val 1823.

**Fig 3.**
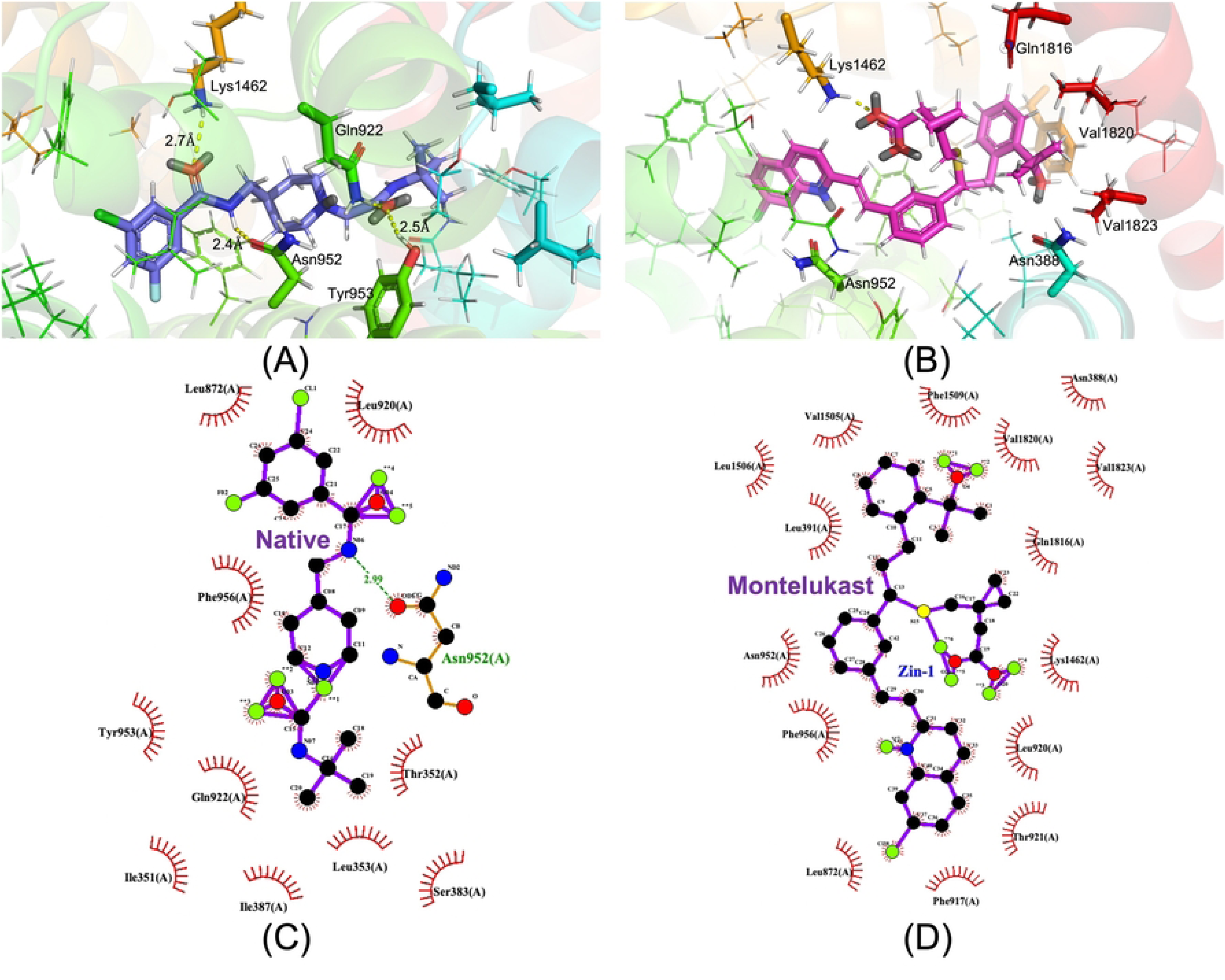
**A, B:** 3D illustration of the docked structures between human Ca_v_3.1 (PDB: 6KZP) and (A) native ligand and (B) montelukast, generated by Pymol. The annotated yellow dotted lines indicate the distance between the atoms, measured in angstrom. **C, D:** 2D illustration of the docked structures between human Ca_v_3.1 (PDB: 6KZP) and (C) native ligand and (D) montelukast, generated by LigPlot+. The green dotted lines represent hydrogen bonding. The red spoked arcs indicate the residues making hydrophobic contacts with the ligand.

Montelukast is a leukotriene receptor antagonist that has been approved by the FDA for the treatment of asthma and allergic rhinitis since 1998 [17]. Millions of asthmatic patients around the world have been taking it regularly, generally as 10 mg once daily by mouth for the prophylaxis of asthma attacks [17]. The side effects of montelukast are minimal and can be tolerated by most patients [17]. The unique chemical structure of montelukast seems to have a low affinity to many other drug targets, and this causes very few known drug-drug interactions [17]. Another advantage of montelukast is its safety in pregnancy [18,19]. This study aims to identify a drug for neurological disorders, such as epileptic seizures, and most current anti-epileptic drugs carry the risk of congenital malformation and adverse prenatal outcomes [20]. Many studies have proven that montelukast does not increase the rate of congenital malformation and can be safely used during pregnancy [18,19].

In addition, montelukast is known to have the ability to cross the blood-brain barrier and could produce neuroprotective effects [21,22]. An animal study showed that montelukast increased the number of neurons by 15% in mice with cranial irradiation [21]. In mice and rats montelukast has been found to reduce neuron loss after a chronic brain injury caused by cerebral ischaemia [22]. A recently published review article also suggested the neuroprotective property of montelukast and its potential use as an anti-epileptic drug, as it reduced the incidence and severity of seizures in epidemiological studies [23]. However, the pharmacological mechanism of these properties is not known. Our docking results show that montelukast may have a high binding affinity to Ca_v_3.1, which could be the reason for its neurological effects. Because of all the above-mentioned characteristics of montelukast, we selected it for further simulations to investigate its binding with Ca_v_3.1.

### Binding free energy calculations

MMPBSA and MMGBSA binding free energy calculations were performed on the docked conformations of both Z944 and montelukast. The results of both approaches show that montelukast binds with a higher affinity than Z944, as shown by the lower binding free energies of montelukast (Table 2). This indicates that the binding complex of montelukast and Ca_v_3.1 is more stable than that of Z944. This increased stability was mainly a result of the van der Waals (vdW) interaction energies (Table 2). The vdW energy difference between montelukast and Z944 is approximately 17 kcal mol^-1^ for both approaches. The change in electrostatic energies (*ΔE_elec_*) is zero for both Z944 and montelukast (Table 2), which could be due to both the ligand and the amino acid residues in the binding site having no significant charges, as *ΔE_elec_*is the long-range interaction between charged atoms.

**Table 2.** Binding free energy calculations between Ca_v_3.1 and its native ligand (Z944) or montelukast by MMPBSA and MMGBSA. *ΔE*_vdW_ is the van der Waals energy, *ΔE_elec_* is the electrostatic energy, *ΔG_SA_* is the solvent accessible surface area energy, *ΔG_PB_* is the Poisson-Boltzmann energy and *ΔG_GB_* is the generalised Born energy. Binding indicates binding free energies. Units of all energies are in kcal mol^-1^.

| Ligand | $\Delta E_{vdW}$ | $\Delta E_{elec}$ | $\Delta G_{SA}$ | $\Delta G_{PB/GB}$ | Binding |
| --- | --- | --- | --- | --- | --- |
| **MMPBSA**<br>Native (Z944) | -40.84 | 0.00 | 0.00 | 22.07 | -23.10 |
| Montelukast | -57.95 | 0.00 | 0.00 | 22.84 | -41.44 |
| **MMGBSA**<br>Native (Z944) | -40.64 | 0.00 | -4.20 | 15.80 | -29.15 |
| Montelukast | -57.86 | 0.00 | -6.08 | 18.59 | -45.57 |

Fig 4 shows the results of the residue decomposition analysis between Ca_v_3.1 and Z944 or montelukast. The residues Phe 956, Leu 920 and Thr921 interacted strongly with both the native ligand and montelukast in both MMPBSA and MMGBSA calculations. The residues Val 1505, Leu 1506, Phe 1509, Val 1820 and Val 1823 interacted with montelukast but not Z944. These interactions could be the causes of the higher binding affinity of montelukast than Z944. In summary, the MMPBSA and MMGBSA simulations indicated that montelukast can form a stable complex with Ca_v_3.1 that has a higher binding affinity than Z944.

**Fig 4.**
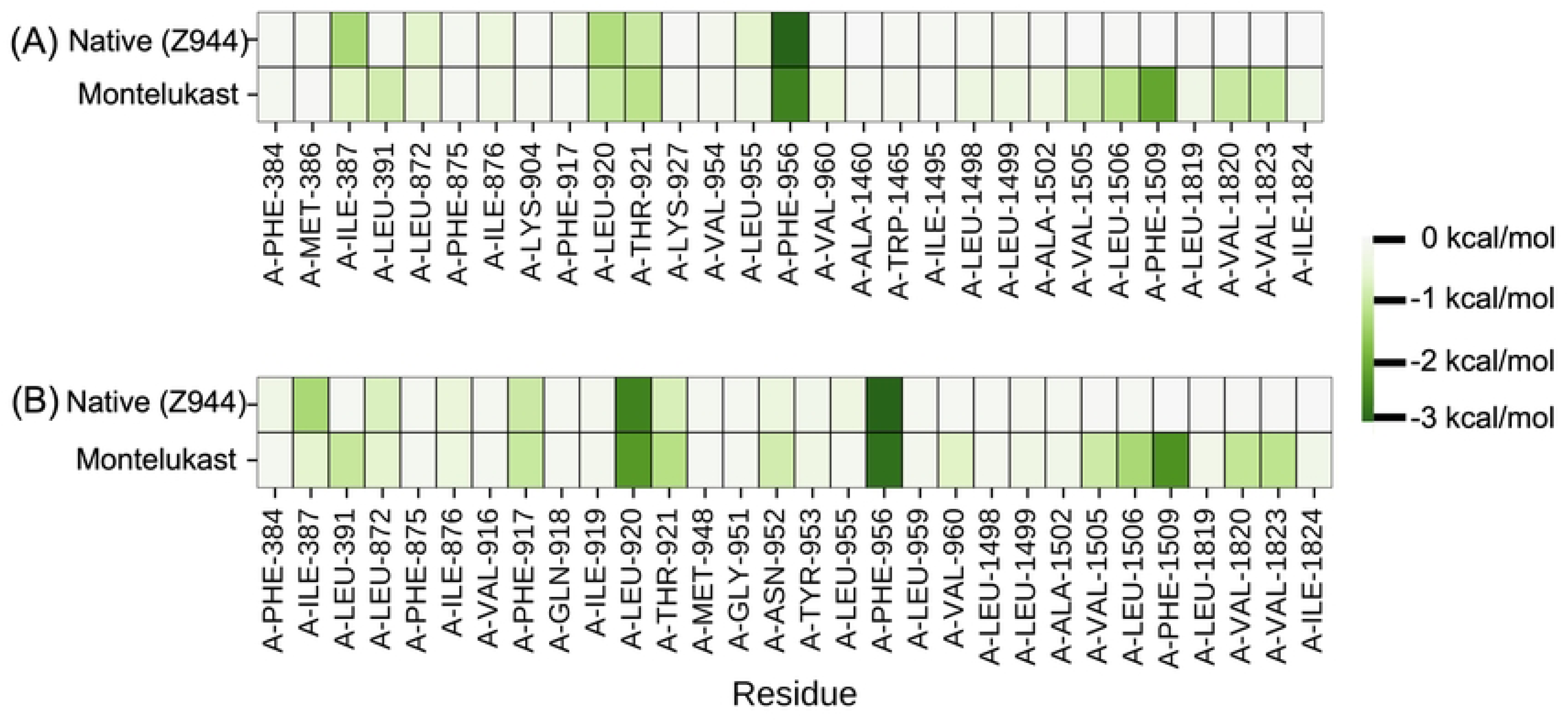
Per-residue decomposition interaction energies (kcal mol^-1^) between Ca_v_3.1 and its native ligand or montelukast, calculated using (A) MMPBSA and (B) MMGBSA methods. Since the native ligand and montelukast bind to distinct residues, panels A and B show different sets of residues. The intensity of the green colour indicates the strength of the energies. Dark green indicates a strong interaction energy.

### Molecular dynamics (MD)

MD was performed on both the crystal structure of the native ligand (Z944) and the best-docked conformation of montelukast with Ca_v_3.1 to investigate their dynamic interactions. Their root mean square deviations (RMSD), root mean square fluctuation (RMSF), radius of gyration (Rg), solvent accessible surface area (SASA), number of hydrogen bonds and interaction energies were calculated.

The RMSD shows the stabilities of the 6KZP-native and 6KZP-montelukast complexes (Fig 5A); their conformations were stabilised by about 2 ns and 17 ns respectively. The 6KZP-native complex was stabilised at an RMSD of about 0.4 nm, whereas that of the 6KZP-montelukast was 0.75 nm. A small RMSD value indicates that there were small differences in the conformation between the docked and MD stabilised structures. In general, an RMSD value of less than 1.0 nm indicates that the docking approach can predict the binding mode of protein-ligand interactions to an acceptable standard [24]. Here, both the stabilised RMSD values of 6KZP-native and 6KZP-montelukast are below 1.0 nm.

**Fig 5.**
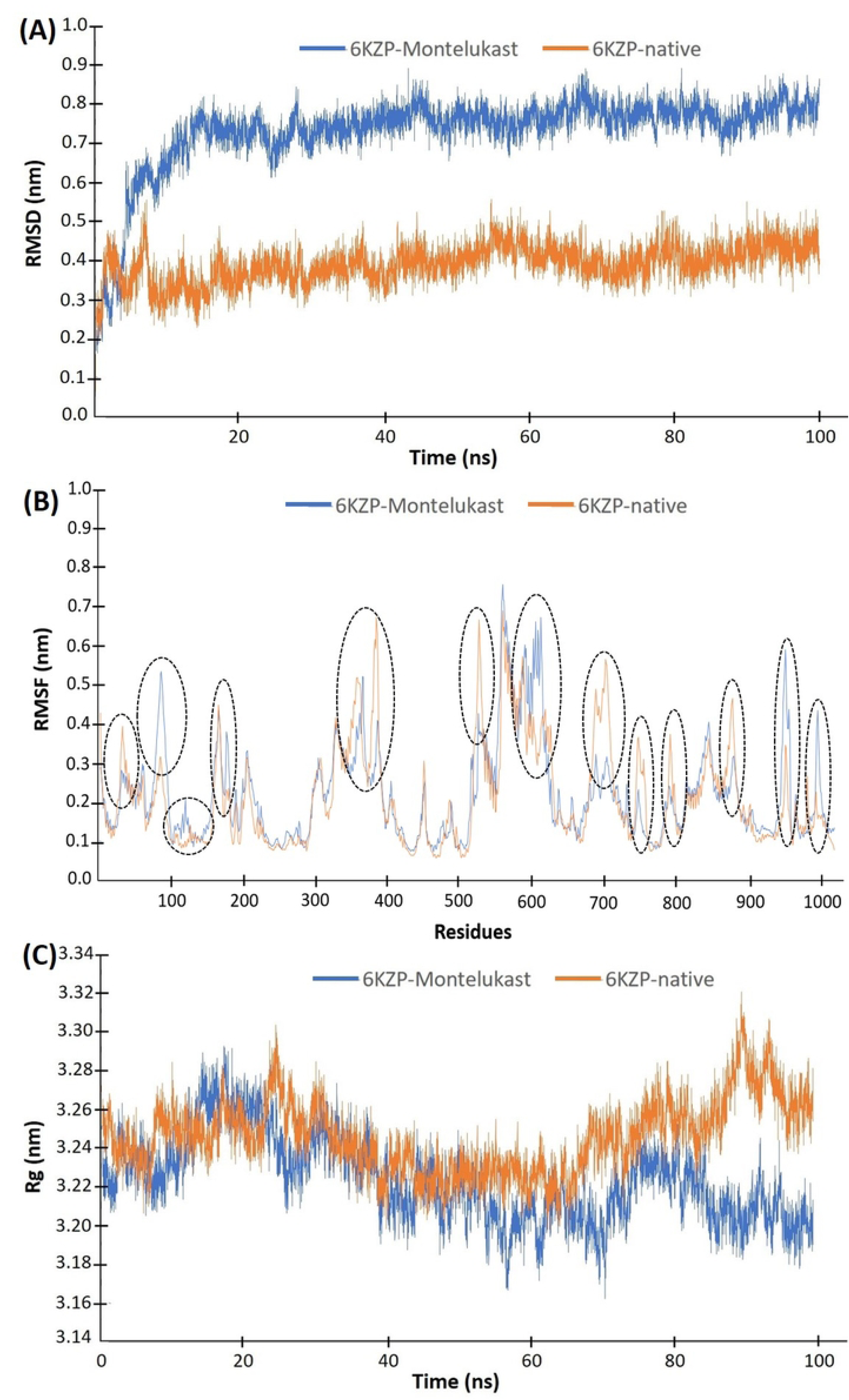
(A) RMSD, (B) RMSF and (C) Rg of the 6KZP-native complex (orange colour) and 6KZP-montelukast complex (blue). The black dotted lines indicate the region with different RMSF between the 6KZP-native and 6KZP-montelukast.

The RMSF results show the fluctuation behaviour of individual residues of the protein (Fig 5B). 6KZP has about 1,200 amino acid residues. The RMSF plots of 6KZP-native and 6KZP-montelukast were similar, indicating that the conformations of both structures have a similar overall variation in stability. The 6KZP-montelukast complex has higher RMSF values than the 6KZP-native complex at residue 85 (Gln 172), 116 (Leu 205), 174 (Asn 263), 593 to 598 (Ser 1334 to Leu 1339), 934 to 935 (Glu 1749 to Thr 1750) and 978 (Asn 1800). These higher RMSF values indicate that these residues are important for the interactions between montelukast and 6KZP, but not for the native ligand. The important interaction residues for the binding between the native ligand and 6KZP are 378 (Val 823), 517 (Arg 1248), 673 to 691 (Cys 1418 to Ser 1436), 735 to 737 (Gln1480 to Pro1482), 741 (His 1486), 777 (Leu 1592) and 860 (Asn 1675).

Rg evaluates the compactness of the complexes (Fig 5C). Over the course of the simulation, the Rg values of the 6KZP-montelukast complex range from 3.17 nm to 3.29 nm, and those of the 6KZP-native range from 3.22 nm to 3.32 nm, suggesting that there may be a small difference in the complexes between the complexes. As shown in Fig 5C, there is a slight increase in the 6KZP-native complex Rg curve and a slight decrease in the 6KZP-montelukast complex after 80 ns of MD simulations. The higher Rg values of the 6KZP-native complex may indicate slightly less compactness, *i.e.*, more flexible structures, than the 6KZP-montelukast complex. The protein of the montelukast complex may be folded slightly more tightly than that of the native.

Hydrogen bonds are one of the strongest types of bonds that are responsible for maintaining the protein structure and provide attractive interaction forces between protein and ligand. In general, hydrogen bonds are stronger than hydrophobic contact and are considered another important facilitator for protein-ligand interactions. Fig 6 shows the number of intra- and inter-hydrogen bonds of the 6KZP-montelukast and 6KZP-native complexes. The intra-protein hydrogen bonds are those within the protein, and the intermolecular hydrogen bonds are those between the ligand and the protein. The number of intra-protein hydrogen bonds was similar for 6KZP-native and 6KZP-montelukast, mainly varying between 720 and 770 (Fig 6A). This level of hydrogen bonds maintained the stabilised structure of the protein. Looking at the intermolecular hydrogen bonds, the 6KZP-native complex tends to have more conformations with two hydrogen bonds than the 6KZP-montelukast (Figs 6B, C and D). This may indicate that the higher binding affinity of 6KZP-montelukast may be caused by other intermolecular forces, such as hydrophobic contact.

**Fig 6.**
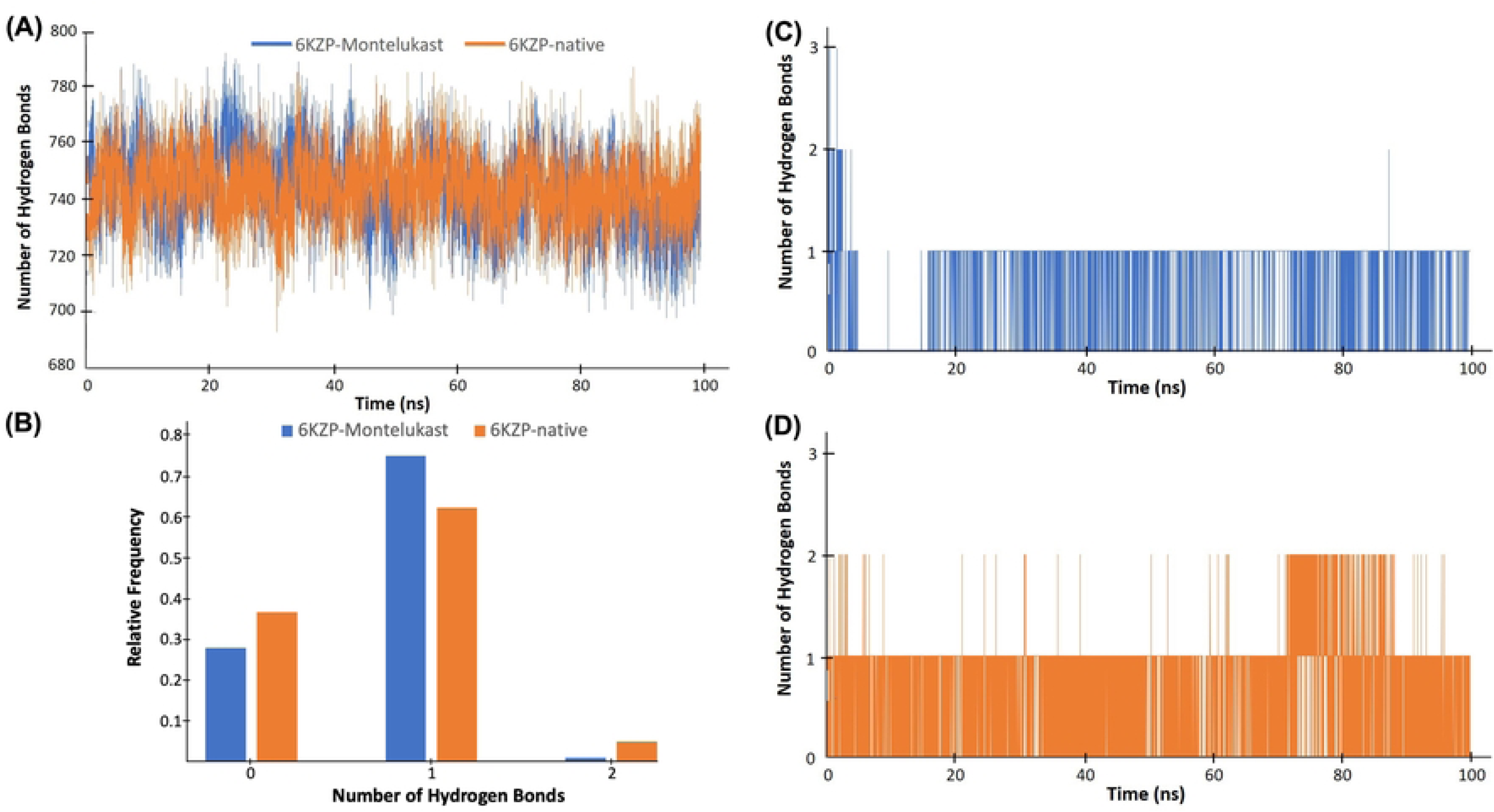
(A) Number of intra-hydrogen bonds for the 6KZP-native complex (orange colour) and 6KZP-montelukast complex (blue). (B) The frequency of the number of inter-hydrogen bonds of the 6KZP-montelukast complex (blue) and 6KZP-native complex (orange colour). (C) Number of inter-hydrogen bonds for the 6KZP-montelukast complex. (D) Number of inter-hydrogen bonds for the 6KZP-native complex.

The 6KZP-montelukast complex has lower solvent accessible surface area (SASA) values than the 6KZP-native complex over the 100 ns MD simulation (Fig 7A). The final SASA values of the 6KZP-montelukast and the 6KZP-native complexes are about 610 nm^2^ and 520 nm^2^, respectively. A high SASA value indicates that a high proportion of the complex is surrounded by solvent molecules, whereas a low SASA value indicates that a large number of solvent molecules are buried inside the complex. Thus, the structure of the 6KZP-montelukast complex has less area covered by solvent molecules and is more compact and probably more stable than the 6KZP-native complex.

**Fig 7.**
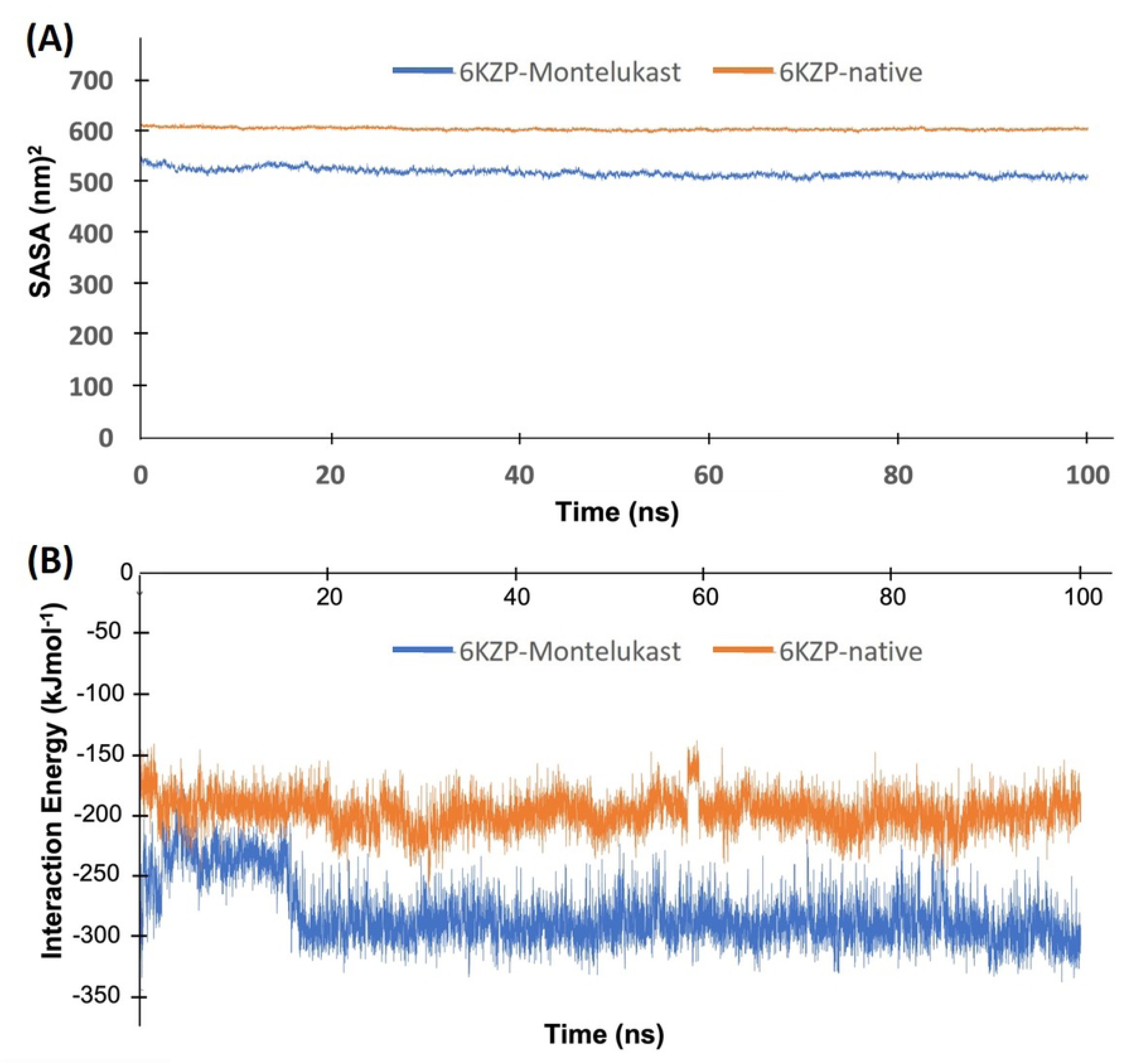
(A) SASA and (B) interaction energies of the 6KZP-native complex (orange colour) and 6KZP-montelukast complex (blue).

The interaction energy calculated by GROMACS is defined as the potential energy of the complex minus the sum of the potential energies of the protein and ligand. A negative interaction energy means that energy is released upon binding; thus, the complex is more stable than the individual protein and ligand. Here, the interaction energy of the 6KZP-montelukast complex stabilised at around −300 kJmol^−1^ after about 20 ns simulation, whereas that of the 6KZP-native complex stabilised at around −200 kJmol^−1^ after about 2 ns (Fig 7B). The lower interaction energy of 6KZP-montelukast indicates a higher binding affinity and stability than the 6KZP-native.

In summary, both the docking and MD simulations indicate a high binding affinity between montelukast and Ca_v_3.1. This supports further study on the use of montelukast as a Ca_v_3.1 inhibitor for the treatment of neurological diseases, such as epileptic seizures.

### Monte Carlo Cell (MCell) modelling

A model utilising MCell and CellBlender was constructed to simulate the interaction between montelukast, calcium ions, Ca_v_3.1, synaptic active membrane, and synaptic vesicles in and around the pre-synapse. It incorporates the opening of Ca_v_3.1 channels near the pre-synaptic bouton, enabling the flow of calcium ions from the extracellular space into the pre-synapse. The influx of calcium ions can trigger the release of neurotransmitter molecules into the synaptic cleft, where they diffuse and initiate signal transmission in the postsynaptic cell. Excessive calcium influx can result in neurological disorders such as epileptic seizures. Conversely, if montelukast blocks the Ca_v_3.1 channels, it can reduce calcium entry and neurotransmitter release, potentially preventing such disorders.

In total, eight 0.2 s MCell simulations were performed using different concentrations of montelukast and different rates of reaction (*k_2_*) between montelukast and the calcium channel (Table 3). The simulations demonstrated that the calcium ions and montelukast competed with the calcium channels. Once the montelukast molecules bound to the calcium channels, they blocked the movement of the calcium ions from the extra-synaptic region to the pre-synapse. This reduced the formation of the complex of calcium ions and synaptic vesicles in the pre-synapse, and thus reduced the amount of neurotransmitter released to the extra-synaptic region.

**Table 3.** The number of calcium ion and vesicle complexes located in the extra-synaptic region at 0.1 and 0.2 seconds in the MCell simulations under different *k_2_* rate constants and initial number of montelukast.

| Number of montelukast molecules | Rate $k_2$ ($M^{-1}s^{-1}$) | Number of the calcium ion and vesicle complex at 0.1 second | Number of the calcium ion and vesicle complex at 0.2 second |
| --- | --- | --- | --- |
| 0 | NA | 2312 | 2665 |
| 500 | $0.5 \times 10^6$ | 2313 | 2654 |
| 500 | $1.0 \times 10^7$ | 2289 | 2671 |
| 500 | $5.0 \times 10^7$ | 2271 | 2588 |
| 500 | $1.0 \times 10^8$ | 2164 | 2539 |
| 1500 | $0.5 \times 10^6$ | 2252 | 2612 |
| 1500 | $1.0 \times 10^7$ | 2239 | 2582 |
| 1500 | $5.0 \times 10^7$ | 1971 | 2262 |
| 1500 | $1.0 \times 10^8$ | 1750 | 1969 |

The results of this study are as expected in our hypothesis. A high concentration of montelukast molecules and high *k_2_* values reduced the number of vesicles docking with the presynaptic membrane and releasing neurotransmitter to the extra-synaptic region. There were about 26% fewer vesicles docking when 1,500 montelukast molecules were presented with a *k_2_*value of 1.0 × 10^8^ M^-1^s^-1^ in the system compared to the absence of montelukast. In contrast, there was almost no change in the number of vesicles docking when comparing the simulation of 500 montelukast molecules with a *k_2_* value of 0.5 × 10^6^ M^-1^s^-1^ and the absence of montelukast (Table 3). When the amount of montelukast was set at 500 and the value of *k_2_*increased from 0.5 × 10^6^ M^-1^s^-1^ to 1.0 × 10^8^ M^-1^s^-1^, there was only a 4% increase in the number of vesicles being released. For the simulations with 1,500 montelukast, the number of vesicles being released increased by 24% when *k_2_* increased from 0.5 × 10^6^ M^-1^s^-1^ to 1.0 × 10^8^ M^-1^s^-1^. Thus, both the concentration of the montelukast and the *k_2_* reaction rate are important factors in these simulations.

Fig 8A and B show the MCell simulation results with the standard parameters, a *k_2_* rate constant of 1.0 × 10^8^ M^-1^s^-1^, with 1,500 montelukast molecules and with no montelukast molecules. The figures indicate that the calcium ions and montelukast compete with each other to bind to the calcium channel located in the extra-synaptic region. After 0.2 s of the simulation, the system was almost equilibrated, and about 700 montelukast molecules were bound to the calcium channels. This reduced the number of calcium ions that could bind to the channels, and thus the number of vesicles being released to the extra-synaptic region.

**Fig 8.**
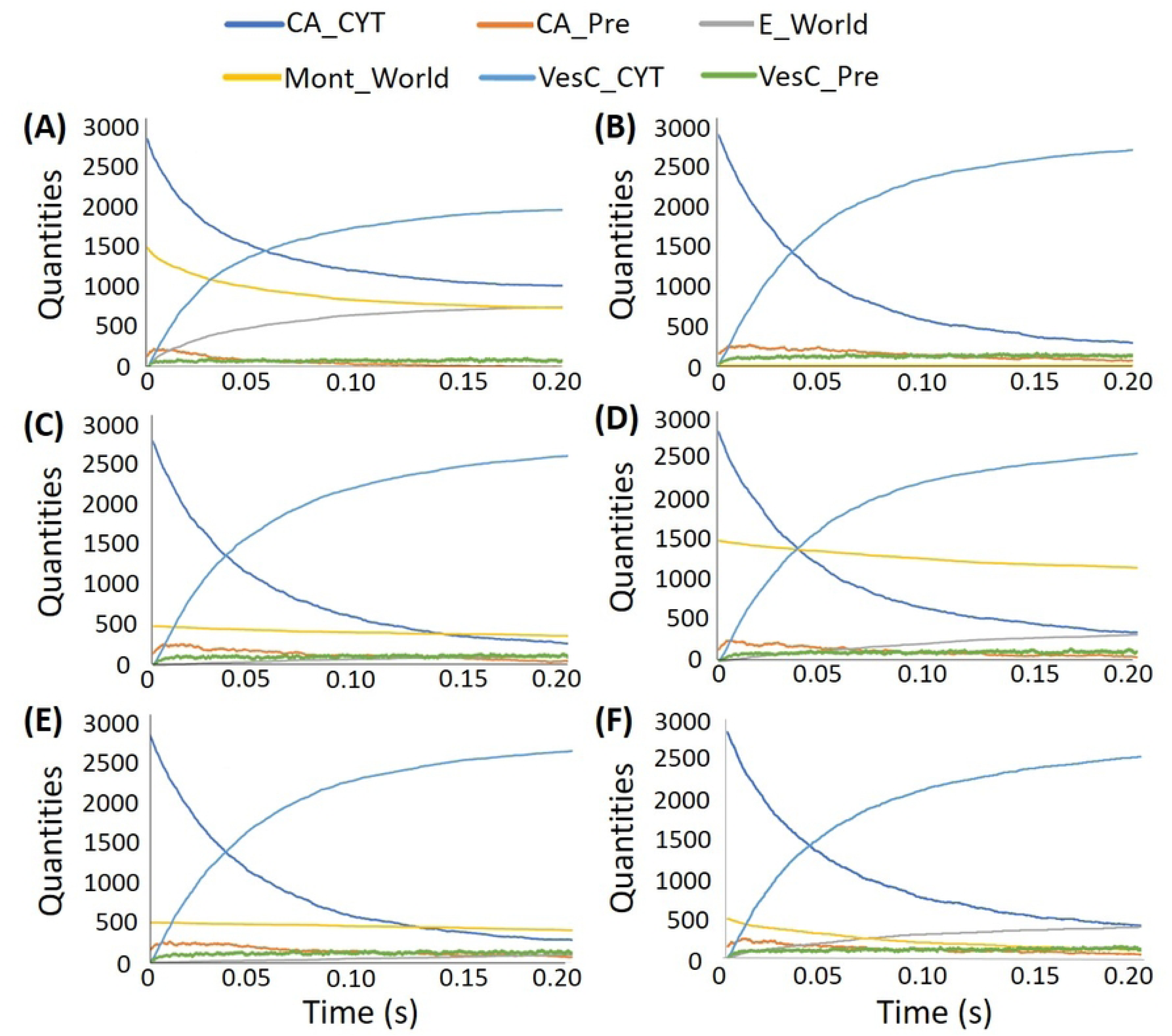
Quantities of the items in the MCell simulations with standard parameters and: (A) 1,500 montelukast molecules and *k_2_* = 1.0 × 10^8^ M^-1^s^-1^; (B) no montelukast molecules and *k_2_* = 1.0 × 10^8^ M^-1^s^-1^; (C) 500 montelukast molecules and *k_2_* = 1.0 × 10^7^ M^-1^s^-1^; (D) 1,500 montelukast molecules and *k_2_*= 1.0 × 10^7^ M^-1^s^-1^; (E) 500 montelukast molecules and *k_2_* = 0.5 × 10^6^ M^-1^s^-1^; (F) 500 montelukast molecules and *k_2_*= 1.0 × 10^8^ M^-1^s^-1^. CA_CYT and CA_Pre are the calcium ions located in the extra-synaptic region and the pre-synapse, respectively, E_World is the complex of montelukast and calcium channel, Mont_World is the montelukast in the whole system, VesC_CYT and VesC_Pre are the complexes of calcium ions and vesicles located in the extra-synaptic region and the pre-synapse, respectively.

Fig 8C and D show the simulations with the standard parameters, a *k_2_* rate constant of 1.0 × 10^7^ M^-1^s^-1^ and with 500 and 1,500 montelukast molecules. Again, the systems seem to be almost equilibrated after 0.2 s. The number of calcium channels blocked by the montelukast increased gradually with time, leading to a reduction in the vesicles being released in the extra-synaptic region. Fig 8E and F illustrate than an increase of *k_2_* rate constant from 0.5 × 10^6^ M^-1^s^-1^ to 1.0 × 10^8^ M^-1^s^-1^ for the system, with a small number of montelukast molecules (500) has a minimal effect on the number of vesicles being released. This may simply be because the lower reaction rate causes the montelukast to be less competitive than the calcium ions in binding to the channels.

## Limitations

Our results indicate that montelukast binding to Ca_v_3.1 is more stable than its native ligand Z944. Montelukast may have the ability to decrease the number of neurotransmitters released from the pre-synapse and potentially has therapeutic effects on neurological disorders, such as epileptic seizures. However, there are several limitations to this study.

Docking and MD can only suggest that montelukast has a high binding affinity to Ca_v_3.1, but cannot indicate whether it is an inhibitor or an inducer. An inhibitor of Ca_v_3.1 may produce the therapeutic effects mentioned in this study. In contrast, an inducer of Ca_v_3.1 may increase the amount of calcium ion flux into the pre-synapse and increase signal transmission between neurons, thus worsening the condition of patients with neurological disorders, such as epileptic seizures. This indistinguishable property is the common drawback of docking and MD. However, most ligands are inhibitors, because induction requires activation of an enzyme, which generally requires conformational changes after binding, and only a very specific ligand structure can achieve this [25]. Thus, there is a high chance that montelukast is an inhibitor of Ca_v_3.1.

Other limitations that may affect the interpretation of the results of this study are the result of uncertain reaction rates and the simplification of the MCell model. The reaction rates of montelukast binding and blocking the calcium channel could not be found in the literature; thus, several different values were explored in this study. Although these rates were based on that of another calcium channel blocker nifedipine [26], they may not be precisely appropriate for montelukast. Another limitation is that the exchange of ions and neurotransmitters within the synapse requires many different types of channels and diffusion mechanisms, and these are far more complicated than our model. There are also many organelles in the human pre-synapse region, and they may affect the diffusion of calcium ions, synaptic vesicles and neurotransmitters. Nevertheless, our MCell results successfully simulated the interaction of montelukast molecules with all other components in the synapse model and showed that montelukast molecules reduced the number of vesicle complexes released from the pre-synapse. Thus, we believe the negative effect of this limitation on the result is minimal.

## Conclusions

Montelukast is an FDA-approved drug that has been used for decades. It has the appropriate pharmacodynamics, pharmacokinetics and pharmacovigilance profile for long-term oral administration [18,20]. Montelukast is considered safe for use in pregnancy, while all others anti-epileptic drugs carry the risk of congenital malformation and adverse prenatal outcomes [20]. This study used molecular docking, MD and MCell simulations to illustrate the potential of montelukast in binding to the Ca_v_3.1 and reducing signal transmission between neurons. The results of this study support further *in vitro* investigations of montelukast as a safe and effective treatment for neurological diseases, such as epileptic seizures. The next step of this study is to perform an *in vitro* inhibition assay on Ca_v_3.1 and confirm the inhibition property of montelukast [27].

## Materials and Methods

### Molecular docking

Molecular docking simulations were employed to calculate the binding affinities between Ca_v_3.1 and the 2,115 FDA-approved drugs of the ZINC database subset [9], aiming to identify a currently-used drug that may inhibit the calcium channel. The 3-dimensional (3D) atomic coordinates of the sole Ca_v_3.1 structure derived from *Homo sapiens* in the Protein Data Bank, a ligand-bound Ca_v_3.1 (PDB code 6KZP), was retrieved [28]. This structure was previously determined by single particle cryogenic electron microscopy with 3.10 Å resolution [28]. The ligand bound to the structure was a Ca_v_3.1 selective antagonist, Z944, which was located in the central cavity of the pore domain (Fig 1) [29]. This structure has been used in several studies for Ca_v_3.1 channel simulations, including one that discovered a novel cyclic peptide as a Ca_v_3.1 channel inhibitor [30,31].

Many docking programs have been developed, each of which has its own set of docking algorithms, scoring functions, and optimization methods, resulting in varying degrees of accuracy when applied to different protein-ligand systems [24]. The docking algorithm is used to search all the potential orientations and conformations of the ligand located in the binding site of the protein. The scoring function is used to calculate the binding energies or binding scores of all the potential orientations and conformations [32]. The automated docking suite GOLD v5.5 [33] was used for all docking simulations in this study. GOLD contains several docking algorithms and scoring functions for users to select. According to a previous study [33,34], the genetic algorithm of GOLD achieved an 80-90% success rate in finding the experimental binding modes of a dataset with 85 complexes, depending on the protocols used. ChemPLP achieved the highest success rate among 20 different scoring functions in a docking power test using a dataset of 195 protein-ligand complexes [35]. This combination of docking algorithm and scoring function has also been used for calcium channel docking simulations [36]. Therefore, in this study, the genetic algorithms with 100% search efficiency and the ChemPLP scoring function [37] of GOLD v5.5 were used for all docking simulations. The docking protocols were set with no early termination, and the ‘slow’ option with high accuracy and the default parameters were used. Atoms within an area of 6 Å of the cognate ligands (Z944) in the X-ray crystallographic structures were set as the binding sites (PDB code 6KZP).

To validate the accuracy of our docking protocols in simulating Ca_v_3.1 complexes, we performed a receiver operating characteristic analysis. It has been used in various studies to evaluate the performance of docking simulations, mainly focusing on their ability to distinguish between the ‘true’ hits and the ‘negatives’ [38–40]. In the receiver operating characteristic analysis, the docking scores were used to plot the true positive rate against the false positive rate at various threshold settings. The true positive rate was calculated using the experimentally determined binding affinities from the Zinc-*in-vivo* (ZIV) database [9]. The area under the curve (AUC) was calculated to indicate the capability of the docking procedure in classifying the true hits from the whole dataset. An AUC value of 1.0 indicates a perfect classification model, and a value of 0.5 indicates a random model with no predictive power. In general, an AUC value of 0.7 or above means that the docking scores have acceptable predictive power in distinguishing between binders and non-binders to the protein [41].

The receiver operating characteristic analysis of this study used the docking scores between Ca_v_3.1 and 14,066 compounds in the ZIV database [9] as the ‘negatives’. These compounds are considered negative since they lack specific information indicating their status as T-type calcium channel inhibitors. Nonetheless, there is a possibility that a small subset of them retains the ability to bind to T-type calcium channels. Additionally, 548 T-type calcium channel inhibitors obtained from the ZINC database as the ‘true’ hits. Thus, the total number of complexes used in this analysis was 14,614. All the compounds of the ZIV database have reported bioactivities in animals, including humans. The result of our analysis, shown in Fig 9, indicates the satisfactory predictive power of our docking method. The high AUC value (0.865) indicates that the method has high sensitivity and specificity.

**Fig 9.**
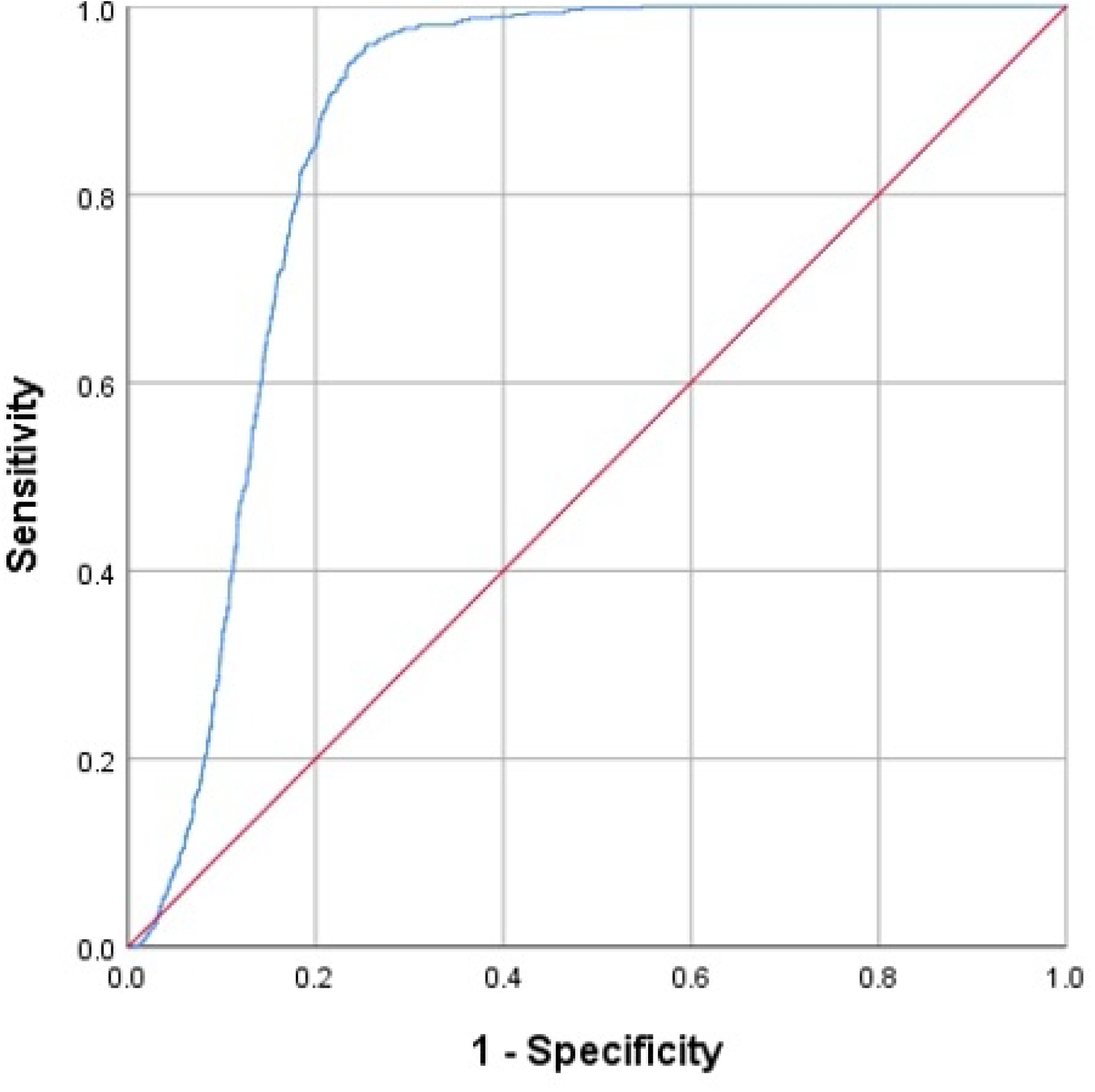
Receiver operating characteristic curves of our docking simulations between Ca_v_3.1 (PDB code 6KZP) and the 14,614 compounds obtained from the ZIV database and ZINC database. The red line is a reference line, which indicates no predictive power, and the blue line is the resulting curve with an AUC value of 0.865.

### Binding free energy calculations

In general, docking may not correctly simulate solvation energy, which is the amount of energy associated with the interactions between the protein-ligand complex and water molecules. This is because such simulations impose relatively high computational demand, and it is not practical to calculate solvation energies while screening a large database. However, solvation energies may significantly affect the rate and magnitude of binding between a protein and a ligand [40]. Thus, in addition to the receiver operating characteristic analysis, molecular mechanics energies combined with the Poisson–Boltzmann (MMPBSA) or generalised Born and surface area continuum solvation (MMGBSA) approaches were performed to calculate the binding free energies of the best-docked conformations of the protein-ligand complex, aiming to study their stability [42].

A study of Sun *et al.* compared the performance of MMPBSA and MMGBSA in estimating binding free energies in 1,800 protein-ligand systems [43]. Both approaches had similar performance: the authors judged that MMGBSA was more useful in ranking individual ligands on multiple proteins, while MMPBSA was more appropriate for ranking multiple ligands on the same protein [43]. It is difficult to select which method is more appropriate for this study, which includes only two ligands (the native and the selected FDA-approved drug) and one protein (Ca_v_3.1). Thus, both methods were used here.

Several software packages have been developed to perform MMPBSA and MMGBSA calculations, such as gmz MMPBSA [44], MMPBSA.py [45] and mm pbsa.pl [46]. They generally require special structural preparation of the proteins and ligands, and they only have command-line interfaces. This study used a recently developed web server, fastDRH, to calculate both the MMPBSA and MMGBSA free energies [47]. It has an interactive online platform and only requires the commonly known PDB format of the protein and ligand as the input structural information. FastDRH has also implemented the most recently developed AMBER (Assisted Model Building and Energy Refinement) protein force field [46], ff19SB [48], and the optimal point charges (OPC) water model. This combination has shown advanced predictive power in modelling protein structures [48] and was therefore used in this study.

### Molecular dynamics (MD)

The aim of performing MD was to validate the docking results and to investigate protein-ligand dynamic interactions. MD simulates the motions and flexibility of a protein-ligand complex over a period of time, rather than a set of conformations as in the docking simulation. Thus, MD is more computationally demanding than docking and generally requires a high-performance computer. In this study, the ECDF Linux Compute Cluster (Eddie) of the University of Edinburgh with GPU acceleration was used to perform 100 ns MD simulations with a 2 ps time step between the Ca_v_3.1 channel (PDB code 6KZP) and Z944 or the selected FDA-approved drug.

In this study, utilities from the GROMACS suite [49] were used for all MD simulations and analyses. We used ‘gmx mdrun’ for MD simulations [49], ‘gmx rms’ to calculate the root mean square deviations (RMSD) of the Ca_v_3.1 backbone associated with ligand-Ca_v_3.1 complex, ‘gmx rmsf’ to calculate the root mean square fluctuation (RMSF) of the protein amino acid residues, ‘gmx gyrate’ for the radius of gyration (Rg), ‘gmx sasa’ to estimate the solvent accessible surface area (SASA) [50], ‘gmx hbond’ to count the number of hydrogen bonds, and ‘gmx energy’ to calculate the interaction energies between the protein and ligands.

The atom types and parameters of both the Ca_v_3.1 channel and the selected drug molecule were generated using the CHARMM36 force field [51]. The TIP3P water model was introduced to the system in a cubic unit cell shape [52]. Anions (*Cl^−^*) were also added to neutralise the overall charge of the system. This ensures proper representation of electrostatic interactions and satisfactory system stability, preventing artificial polarisation, water distribution imbalance, and long-range interactions. This is a conventional practice in MD for more realistic and reliable simulations of protein-ligand complexes. Energy minimisation was performed on the system, to determine a molecular arrangement of all atoms that avoids steric clashes, using the steepest descent minimisation with 5×10^4^ steps. After minimisation of the solvated and electroneutral system, equilibration was performed to ensure that the solvent and ions around the protein-ligand system have the appropriate molecular geometry at a suitable pressure, volume and temperature [53]. In general, equilibration contains two phases: the constant number of particles, volume and temperature (NVT) equilibration and the constant number of particles, pressure and temperature (NPT) equilibration. In this study, both NVT and NPT were performed for 5 ns with a 0.2 ps time step, while the protein and ligand were positionally restrained. The Berendsen temperature coupling method and Parrinello-Rahman barostat pressure coupling method were used to maintain the system at 300K and 1 bar [49]. These techniques approximate the physiological temperature and atmospheric pressure for realistic conditions for the simulation [49]. The Lennard Jones potential was used to estimate the short-range interaction with a cut-off value of 12 Å. The long-range interactions were calculated by the Particle Mesh Ewald (PME) method, which has shown to have an appropriate balance between computational cost and scientific reliability [54].

### Monte Carlo Cell (MCell) modelling

After performing docking and MD to determine the potential Ca_v_3.1 inhibitor from the 2,115 FDA-approved drugs, MCell modelling was conducted to simulate the interactions between the selected drug, calcium ions, Ca_v_3.1, synaptic active membrane and synaptic vesicles in and around the pre-synapse. In the human nervous system, when an action potential propagates into an axon, the Ca_v_3.1 channels close to the pre-synaptic bouton open and calcium ions from the extracellular space flow into the pre-synapse down the concentration gradient of calcium from the extra- to intra-presynapse. The calcium ions in the pre-synapse then trigger the synaptic vesicle to fuse with the pre-synaptic active membrane, and the neurotransmitter molecules in the synaptic vesicle are released into the synaptic cleft, where they diffuse, with some reaching postsynaptic receptors, initiating various forms of signal transmission in the postsynaptic cell. In particular, binding to iontropic receptors such as AMPA or NMDA receptors leads to influx of charge carried by ions into the postsynaptic cell. If the magnitude of this influx is too high, neurological disorders, such as epileptic seizures, may occur [1]. Blocking the Ca_v_3.1 channel by inhibitors may reduce the amount of calcium influx into the pre-synaptic region, and decrease the number of neurotransmitter molecules being released. The resultant reduction in excitation may alleviate the severity of symptoms associated with neurological disorders.

This study employed an open-source 3D graphical software, CellBlender bundle version 4.0.5 with Blender 2.93, to create and visualise the synapse model. Blender is a software toolset that has been used to generate animated films, video editing, fluid dynamic simulations and more [55]. Thousands of add-ons have been developed for different purposes, including CellBlender, which is embedded with MCell (version 4.0.5) [56–58]. MCell is a comprehensive modelling environment that uses Monte Carlo Cell simulations to investigate the characteristics and geometries of particles and their reactions, including synaptic plasticity in dendritic spines [59–61].

The synapse model created in this study contains three parts: the pre-synapse, post-synapse and extracellular regions (Fig 10). The volume and surface area of the pre-synapse region were 0.21 µm^3^ and 2.27 µm^2^, respectively; those of the post-synapse region were 0.15 µm^3^ and 2.13 µm^2^, and those of the extracellular region were 4.63 µm^3^ and 16.04 µm^2^, respectively [62]. Calcium channels were placed on the upper surface of the pre-synapse, with a density of 104 per µm^2^ (Fig 10C) [62]. The bottom surface of the pre-synapse was assigned as an active membrane for the release of synaptic vesicles (Fig 10D). These values were adopted from a recent study [62], which used MCell simulations to investigate the inhibition effects of gadolinium ions on calcium channels located on the pre-synapse.

**Fig 10.**
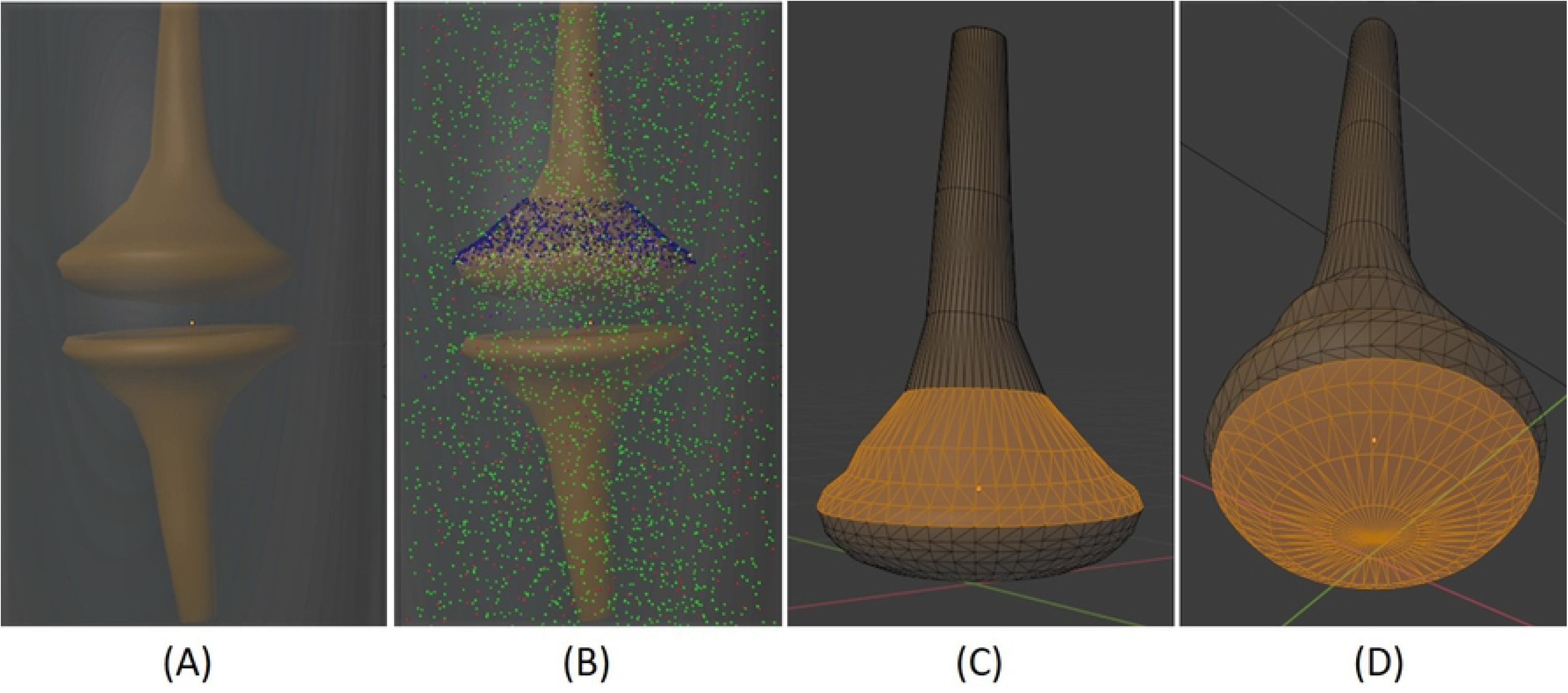
Model of synapse: (A) The pre-synapse (top) and post-synapse (bottom); (B) The synapse model with calcium ions (green), montelukast (red), calcium channels (blue) and calcium pump (black); (C) The pre-synapse region (orange) where the calcium channels were located; (D) The pre-synapse region (orange) of the active membrane where the vesicle and calcium ion complexes were released.

Four reactions were used to describe the interactions between the selected FDA-approved drug, calcium ions and the synaptic vesicles in different compartments:

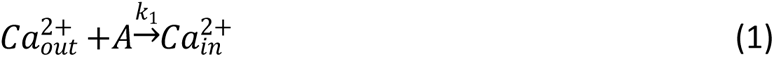

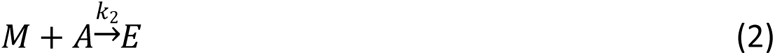

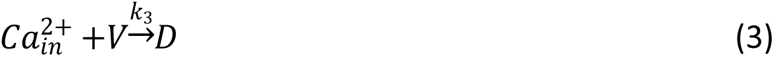

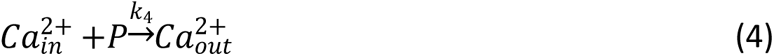

where 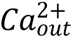 and 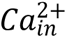 indicate the calcium ions located outside and inside the pre-synapse, respectively. The calcium channel (A) allows calcium ions to travel into the pre-synapse. Thus, equation 1 indicates the diffusion of calcium ions from the extracellular region into the pre-synapse through the calcium channels. Equation 2 indicates the selected FDA-approved drug M blocking the calcium channel (A) and forming a complex of the drug and calcium channel (E). V and D in equation 3 are the vesicles located inside the pre-synapse and the calcium ion and vesicle complex, respectively. In human pre-synapse, calcium ions are pumped out of the neurons by several mechanisms, such as ATP-driven pumps and Na^+^/Ca^2+^ exchangers [63]. This study collectively calls all mechanisms that remove the calcium ions ‘calcium pumps’. The P of equation 4 is the calcium pump that moved calcium ions out of the pre-synapse to the extracellular region.

The rate of calcium flux into the pre-synapse after binding to the calcium channel is denoted *k_1_*. The rate that the selected drug molecules bind and block the calcium channels is denoted *k_2_*. The rate that calcium ions bind to the vesicles inside the pre-synapse is denoted *k_3_*. The rate of the calcium ions being pumped out of the pre-synapse is denoted *k_4_*. The values of these rate constants (except *k_2_*) were adopted from the study by Sutresno *et al*. [62] and are listed in Table 4. As the rate constant *k_2_* could not be found in the literature, we used four different values to explore the effects of the selected drug. According to the study by Mery *et al.* [26], the reaction rate constant of the calcium channel inhibitor nifedipine ranges from 1 × 10^6^ to 4.47 × 10^6^ M^-1^s^-1^. The docking results of this study (Table 1) show that the docking scores of the selected drug and nifedipine were 100.71 and 47.78, respectively. Thus, we believe the reaction rate of the drug is higher than that of nifedipine, *i.e.*, at least 4.47 × 10^6^ M^-1^s^-1^. We explored the effects of four different *k_2_* values: 5 × 10^6^ M^-1^s^-1^, 1 × 10^7^ M^-1^s^-1^, 5 × 10^7^ M^-1^s^-1^ and 1 × 10^8^ M^-1^s^-1^ (Table 4).

**Table 4.**
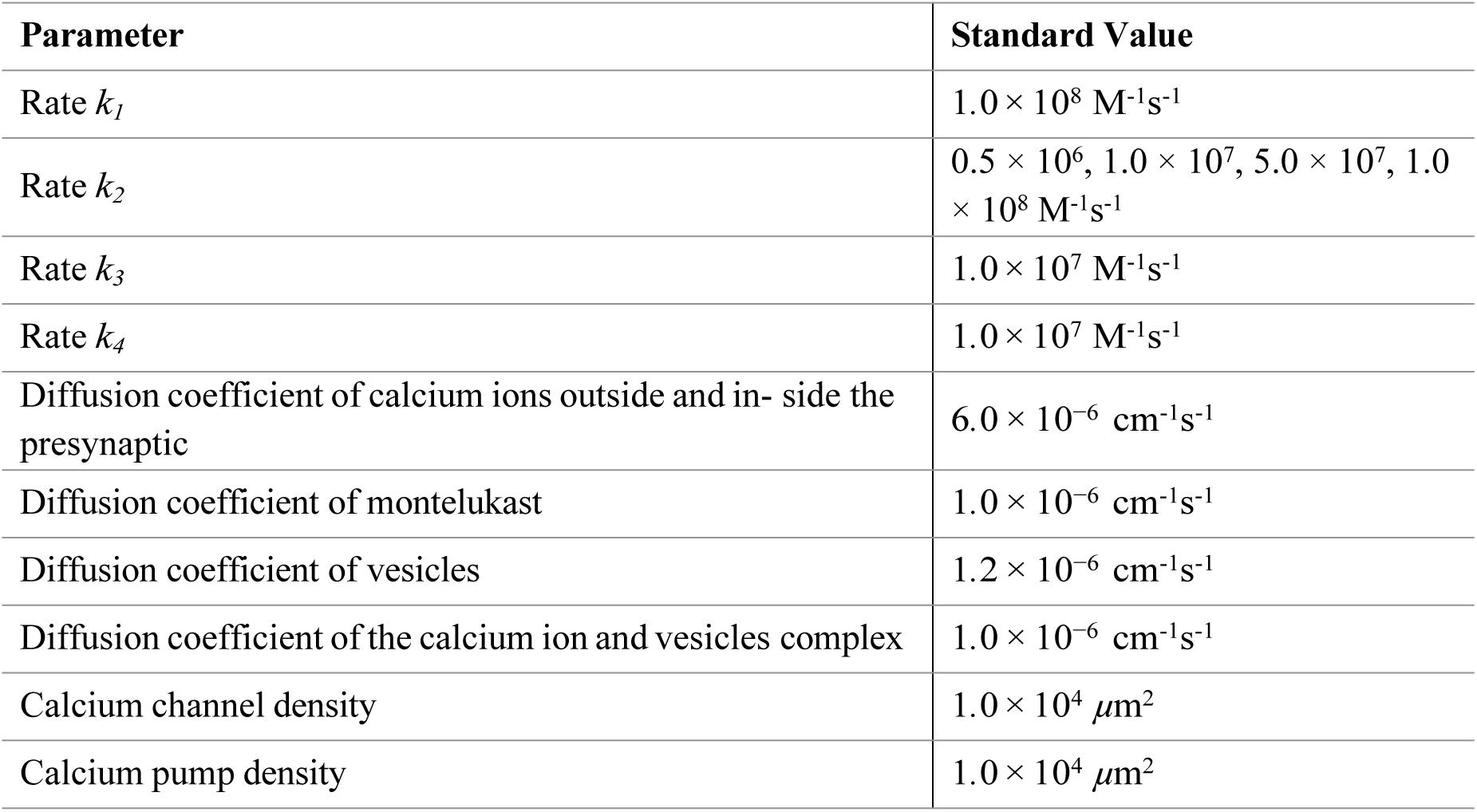
Standard value of parameters in the MCell simulations.

| Parameter | Standard Value |
| --- | --- |
| Rate $k_1$ | $1.0 \times 10^8 \text{ M}^{-1}\text{s}^{-1}$ |
| Rate $k_2$ | $0.5 \times 10^6, 1.0 \times 10^7, 5.0 \times 10^7, 1.0 \times 10^8 \text{ M}^{-1}\text{s}^{-1}$ |
| Rate $k_3$ | $1.0 \times 10^7 \text{ M}^{-1}\text{s}^{-1}$ |
| Rate $k_4$ | $1.0 \times 10^7 \text{ M}^{-1}\text{s}^{-1}$ |
| Diffusion coefficient of calcium ions outside and in- side the presynaptic | $6.0 \times 10^{-6} \text{ cm}^{-1}\text{s}^{-1}$ |
| Diffusion coefficient of montelukast | $1.0 \times 10^{-6} \text{ cm}^{-1}\text{s}^{-1}$ |
| Diffusion coefficient of vesicles | $1.2 \times 10^{-6} \text{ cm}^{-1}\text{s}^{-1}$ |
| Diffusion coefficient of the calcium ion and vesicles complex | $1.0 \times 10^{-6} \text{ cm}^{-1}\text{s}^{-1}$ |
| Calcium channel density | $1.0 \times 10^4 \mu\text{m}^2$ |
| Calcium pump density | $1.0 \times 10^4 \mu\text{m}^2$ |

In addition to the rate constant, the diffusion coefficient is also an important factor in our simulations. The diffusion coefficient indicates the distance that a molecule diffuses in the medium per unit of time. The diffusion coefficient values used in this study (Table 4) were adopted from a previous study [62], which simulated the interaction between calcium ions and gadolinium ions within calcium channels that regulate entry into the pre-synapse. Two different ratios of the number of calcium ions and the selected drug were used: 6:1 (*N* = 3000 and 500) and 2:1 (*N* = 3000 and 1500). Again, these ratios were adopted from the study by Sutresno *et al.* [62]. These ratios are a rough estimation because no such data can be found in the literature.

The simulation protocol of this study is also similar to that of Sutresno *et al.* [62]: all the calcium channels were in their open states, and they were ready for binding to either the calcium ion or the FDA-approved drug molecule. As no information can be found in the literature with regard to how long the drug molecule could bind to the calcium channel, we simulated that neither the calcium ion nor the drug molecule would unbind from the calcium channel once binding occurs. We initialised the system with specified numbers of calcium ions and drug molecules evenly distributed throughout the extra-synaptic region. The drug molecules competed for the calcium channels, until they were all occupied In summary, this MCell model illustrates the effect of the selected drug on the inhibition of calcium channels. Its effect on the number of calcium ions in different compartments and the number of vesicles being released into the synaptic cleft under different reaction rate constants, diffusion constants and number of drug molecules were also demonstrated.

## Acknowledgement

We are grateful to the National Center for Multiscale Modeling of Biological Systems (MMBios) of the University of Pittsburgh for teaching the authors to build MCell models.

## Author Contributions

Conceptualization: Pedro Fong, David Sterratt, Melanie Stefan, Susana Roman Garcia.

Data curation: Pedro Fong.

Formal analysis: Pedro Fong, David Sterratt, Melanie Stefan.

Investigation: Pedro Fong.

Methodology: Pedro Fong, David Sterratt, Melanie Stefan.

Project administration: David Sterratt, Melanie Stefan, Susana Roman Garcia.

Resources: David Sterratt, Melanie Stefan.

Software: David Sterratt, Melanie Stefan.

Supervision: David Sterratt, Melanie Stefan, Susana Roman Garcia.

Validation: David Sterratt, Melanie Stefan, Susana Roman Garcia.

Visualization: Pedro Fong, David Sterratt, Melanie Stefan.

Writing – original draft: Pedro Fong.

Writing – review & editing: David Sterratt, Melanie Stefan, Susana Roman Garcia.

## Disclosure statement

No potential conflict of interest was reported by the author(s).

## Notes

### Competing Interest Statement

The authors have declared no competing interest.

